# Enabling Success Through Transformative Practices in STEM: The Effects of Applying an Anti-Deficit Framework to Diversity and Equity Programming

**DOI:** 10.1101/2023.06.22.546152

**Authors:** D’Anne S Duncan, Jhia LN Jackson, Sally Collins, Arianne Teherani

## Abstract

Diversity, equity, and inclusion (DEI) programming and literature for historically underrepresented and marginalized students in science, technology, engineering, and mathematics degree programs often focuses on illuminating the challenges they face. The repeated emphasis on negative experiences creates a deficit-focused thread of inquiry that may unintentionally reinscribe persistent disparities and inequities. In this practice brief, we describe the positive effect of adapting anti-deficit framework with social career cognitive theory in developing and evaluating an Initiative for Maximizing Student Development program at a biomedical sciences graduate school, as well as how other institutions can explore, implement, and evaluate transformative DEI practices. In identifying and emphasizing the enablers of success while facilitating structured opportunities for personal and professional identity development, students and program leadership align values and goals to increase academic and scientific development, as well as community and social support.

---

The overarching discourse on diversity and equity in higher education focuses extensively on the lack of historically underrepresented and marginalized (HUM) students in science, technology, engineering, and mathematics (STEM) careers and the difficulties HUM students encounter when they pursue higher STEM degrees (Carlone & Johnson, 2007; Gardner 2008; Gibau 2015; Tull et al., 2012).^1^ The repeated emphasis on the negative experiences and unique difficulties encountered by these students has created a deficit focused thread of inquiry in which research continually identifies institutional mismatches and achievement gaps as compared to their non-HUM peers. However, identifying deficits does not account for the positive experiences and successes of these students, nor does it acknowledge that there are many HUM students who select and successfully complete higher STEM degrees, pursue STEM careers, or choose to use their STEM degrees in creative ways. Little research has focused on understanding the educational experiences and factors that examine and support HUM students’ success during their graduate STEM education. How we support and encourage HUM students to succeed as they enter higher education institutions is particularly important as the early years of graduate education play a formative and vital role in career decision making (Fuhrmann, 2016; Williams et al., 2017).

Programmatic efforts to date have focused on promoting HUM student success in their studies, degree completion, and STEM career choice. Evidence in graduate education has pointed to the effectiveness of academic and social support programs that help HUM students successfully navigate the graduate school experience (Gibau, 2015; Peteet & Lige, 2015; Salto et al., 2014; Tull et al., 2012; Wilson et al., 2018). These programs have included social support through measures such as community building and faculty mentoring (Tull et al., 2012; Wilson et al., 2018), academic support through interventions focused on outcomes such as candidacy examination and dissertation preparation (Tull et al., 2012; Wilson et al., 2018) or the provision of faculty mentors, and professional development support through career and identity exploration.

We understand little about the evolving contextual individual and program facets that support HUM graduate student success during graduate training. Although evidence exists on the success of supportive diversity and equity programs, this work often draws from and inadvertently contributes to the deficit focused thread of inquiry which may unintentionally reinscribe persistent issues rather than moving forward with the changing values and practices of HUM students. A robust understanding of these facets and how they interact over time to create supportive experiences will allow new and existing programs to efficiently develop or refine their support structures while guided by a framework that attends to contemporary forms of success for HUM students. Therefore, we sought to characterize, utilizing social career cognitive theory through the lens of the anti-deficit framework, how a program and its individual components developed to support HUM graduate students in the biomedical sciences creates academic and social support.

## Theoretical Framework

Social career cognitive theory (SCCT) recognizes professional development as a process involving simultaneous career and identity development (Lent et al., 1994; Lent et al., 2018). The theory highlights three factors that influence each of these facets of SCCT: *socio-cognitive* mechanisms as in the student’s belief system, *person* factors as in the student’s personal characteristics and aptitude, and *contextual* variables as in the features of the environment or culture that promote or inhibit choices (Lent et al., 1994). Existing studies of HUM student graduate training and career choices have examined socio-cognitive mechanisms (Carlone & Johnson, 2007; Gardner 2008; Gibau 2015), person factors (Carlone & Johnson, 2007; Gardner 2008; Gibau 2015), and to a lesser extent, contextual variables (Mills et al., 2019). The anti-deficit framework can enhance SCCT by orienting the line of inquiry about the facets of the theory towards the enablers of persistence and success, rather than the ongoing structures that perpetuate persistent disparities and inequities in achievement (Harper, 2010). Higher education institutions are in a vital position to account for shifting socio-cognitive mechanisms and person factors while evaluating and shaping contextual variables to create opportunities that promote multifaceted career development for HUM students.

## Research Methods

Our study was a qualitative constructivist grounded theory study of HUM graduate students participating in an early-graduate school fellowship (Charmaz 2014). Constructivism recognizes that meaning is socially constructed and will differ based on who is constructing meaning, even when considering the same phenomenon. Our study was conducted at the University of California, San Francisco (UCSF) which is the graduate-only, health and life sciences-focused campus of the 10-campus University of California system. The UCSF Graduate Division degree programs are within all professional schools of Medicine, Dentistry, Nursing, and Pharmacy and Global Health Sciences at UCSF. The Institutional Review Board of the University of California approved our study as exempt.

Participants were three cohorts of IMSD fellows between 2016-2018. Each year, six HUM students were^2^? To understand the systems that support success during that early graduate student trajectory, we invited all students to participate in the interviews once at the middle of the fellowship (end of year 1) and once at the end (end of year 2). Fellows were sent up to three reminders to participate. Interviews were conducted either by the evaluator of the program or trained research analysts and lasted between 15 and 45 minutes. We analyzed the data in an iterative fashion as we collected data using a constant comparative approach during the interview (Corbin & Strauss 2008).

The interview guide was developed with a focus on the contextual factors’ component of the SCCT framework, as well as the anti-deficit focus on success and achievement. We pilot tested the protocol with two IMSD students who had completed the program and refined it for clarity.

The final questions asked fellows about:

- overall experiences in graduate school and during the IMSD fellowship;
- experiences within and perception of the IMSD fellowship and program;
- value, if any, of the IMSD activities and meetings to students’ academic and social achievement;
- whether and how the program increased opportunities available to students;
- individuals who provided the student with guidance during their education and what forms the guidance assumed; and
- overall career goals and impact of program, if any, on career goals (end of program only).

### Data Analysis

We used a constant comparative approach during the interviews to collect and analyze data in an iterative fashion (Corbin & Strauss 2008). In the first two years of interviews, we identified and explored themes in depth during subsequent interviews with a particular focus on how the IMSD experiences supported success and achievement. In the first year of data collection, two investigators analyzed 2 interviews to create the initial codebook and refined the interview guide to include more probing questions. The initial codebook was applied to two additional interviews by both investigators and refined further. By the end of the first year of data collection, we had a final codebook which we changed minimally in the latter two years of data collection. The final codebook was applied to all interviews and the investigators reviewed each other’s code to make sure they agreed. We achieved theoretical sufficiency by the second interview in the third year of data collection when we noticed no new patterns and codes merging from the data (Charmaz 2014). We decided to conduct all remaining interviews to complete the third year of data collection.

### Researcher Reflexivity

The research team consisted of a medical education researcher who was the evaluator for the IMSD program, a research analyst, an IMSD Co-PI, and a graduate student researcher. The evaluator and research analyst jointly conducted interviews and analyzed the data. The evaluator is a faculty member in the School of Medicine and an equity and identity researcher with no administrative, teaching, or assessment role in the IMSD program or Graduate Division. The research analyst, also in the School of Medicine, had no relationship to the IMSD program. The IMSD Co-PI oversaw the programmatic elements, served as the primary point of contact for the IMSD students, and supported data analysis and implementation of findings. The graduate student researcher, a social science doctoral candidate who was not eligible for participation in IMSD, also supported the analysis and implementation of findings. All authors provided their ongoing perspectives on the interpretation and write up of the results, particularly as they represent diverse groups of stakeholders within the institution. Collectively, these experiences may have influenced the interpretation of data. However, the research team came from diverse expertise and backgrounds which mitigated potential bias and created a robust final analysis and synthesis.

## How the Use of Social Career Cognitive Theory and the Anti-Deficit Framework Can Lead to Transformative Practices

In this article, we characterize the core and individual features of UCSF’s IMSD Program, its impact on creating academic and social support for HUM graduate students in the biomedical sciences, and how the guidance of SCCT and Harper’s anti-deficit achievement framework contributed to the program’s successes. Particularly, our commitment to utilizing an anti-deficit framework created an underlying shared value that influenced our work, interpersonal interactions, and program evaluation.

SCCT is a framework for interpreting the development of career interests, the academic and career decision-making processes, and the performance and persistence of graduate students in these pursuits (Lent, Brown, Hackett,1994: 2002). This framework addresses research gaps on ethnicity and career self-efficacy by emphasizing personal agency and extra-personal factors that support or constrain agency, allowing for an accounting of systemic social inequities as ongoing processes that create structural opportunities and barriers (Hackett & Byars, 1996). Thus, to promote strong, pragmatic, and positive academic and career self-efficacy beliefs, students need to be supported in their personal identity development along with their academic and career development, while explicitly acknowledging the ways that experiences of racism, sexism, and similar discriminatory processes of social inequity are influencing their efficacy and outcome expectations (Hackett & Byars, 1996).

We complemented our use of SCCT with Harper’s anti-deficit achievement framework (Harper, 2010). Harper argues that there is a plethora of descriptive research on these issues and while such research has value, it is not particularly useful for faculty or administrators who are interested in adapting their policies, practices, and programs to support the enablers of persistence and success for the HUM students, thereby increasing representation, retention, and achievement. When used in conjunction with existing theories, it prompts researchers to “ask why is _ happening instead of the already well documented _”, moving research and practices forward in a manner that has short-term and long-term benefits (Harper 2010). The anti-deficit framework was developed to complement existing psychosocial theories by reorienting inquiry that seeks to address persistent representation and achievement inequities towards the enablers of success, rather than the constrainers of choice or factors of failure (Harper 2010).

As shown in Figure 1, the program was developed to include transformative practices that address all aspects of SCCT while utilizing an anti-deficit framework. When working with program leaders and mentors to develop content and interactional practices, we continuously asked guiding questions that deployed the anti-deficit framework. This nuance allowed our program to be refined not only for our specific institution and community of learners, but for the specific strategies of success that they and others before them were using. This way, the program leaders could adapt to and support each unique cohort and account for social and environmental changes, while also encouraging other UCSF community leaders to shift specific practices in targeted, tangible ways that would engage our program participants and make them feel welcomed, valued, and encouraged.

**Figure 1.**
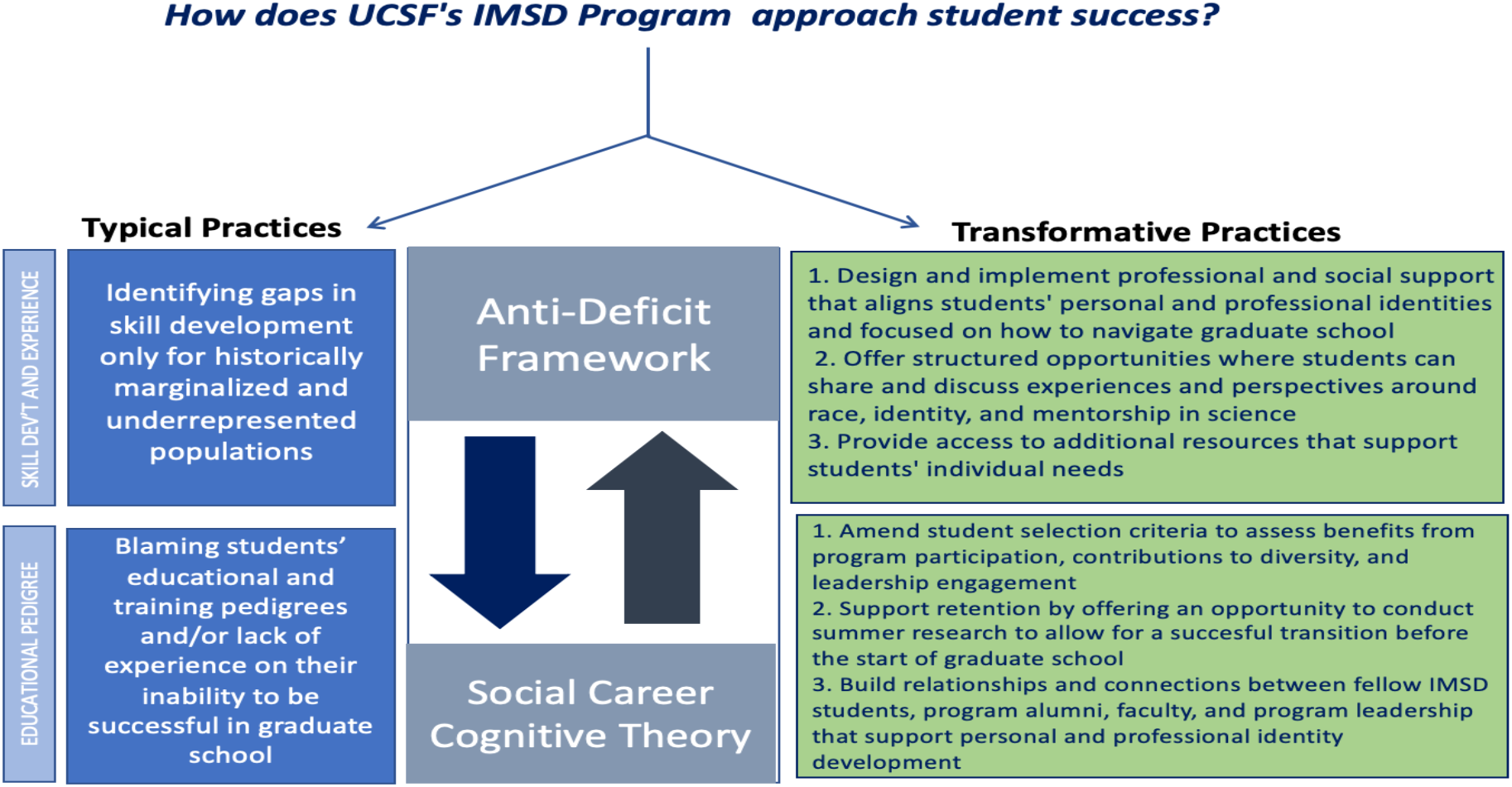
Applying an Anti-Deficit Framework for Transformative Practices

## An Overview of UCSF’s IMSD Program Structure

The National Institute for General Medical Sciences (NIGMS) Initiative for Maximizing Student Development (IMSD) fellowship at UCSF is designed to support the academic and research competitiveness of historically underrepresented and marginalized students (HUMs) in basic science PhD graduate programs and to facilitate their progress toward careers in biomedical research. Since its inception in 1997, the fellowship has supported selected students in the first two years of their graduate studies. Selected students are nominated by their graduate programs and selected by the IMSD Co-PIs, based on the benefits of IMSD on a student’s PhD experience, and their contributions to diversity and leadership engagement experience. The IMSD students participate in 10 PhD programs at UCSF: bioengineering, biological and medical informatics, biophysics, chemistry and chemical biology, developmental and stem cell biology, neuroscience, oral and craniofacial sciences, pharmaceutical sciences and pharmacogenomics, and the Tetrad program (biochemistry, molecular biology, cell biology, and genetics). IMSD program leadership is comprised of two Co-PIs, a tenure-track faculty member and a faculty member who oversees the diversity, equity, and inclusion programs and initiatives. The Co-PIs serve as the primary point of contacts for the IMSD students, lead the professional development and community building activities, and provide extensive mentorship on how to thrive and succeed in graduate school, due to their extensive knowledge and experience in biomedical science research education and training, and strong backgrounds in directly mentoring and supporting HUM graduate students in biomedical science disciplines. The program is also supported by a coordinator who helps with execution of the program’s professional development and community building activities and participates in the evaluation process.

UCSF IMSD students complete two years of codified programming that is evaluated annually and updated to account for contemporary social factors and are subsequently invited to participate in community building and leadership experiences throughout the rest of their academic tenure at UCSF. The fellowship includes the following components: (1) participation in a summer research rotation in which they spend time in an initial lab of their choosing with a principal investigator (PI); (2) rotations of 2-3 months’ duration in three different research labs before joining a lab for their dissertation research by the start of their second year; (3) fostering a community of fellows through ongoing meetings and social events; and (4) monthly seminars and workshops on key topics focused on learning research skills, navigating expected graduate school experiences, career exploration, professional development, and strategies for effective time and project management (see Table 1). Collectively, the combination of these workshops and community building experiences are designed to create affirming environments that celebrate and uplift students’ personal identities and lived experiences while discussing race, identity, mental health, and well-being within the context of personal and professional development. The following subsections detail some of the results from this program development, evaluation, and implementation approach.

**Table 1.**
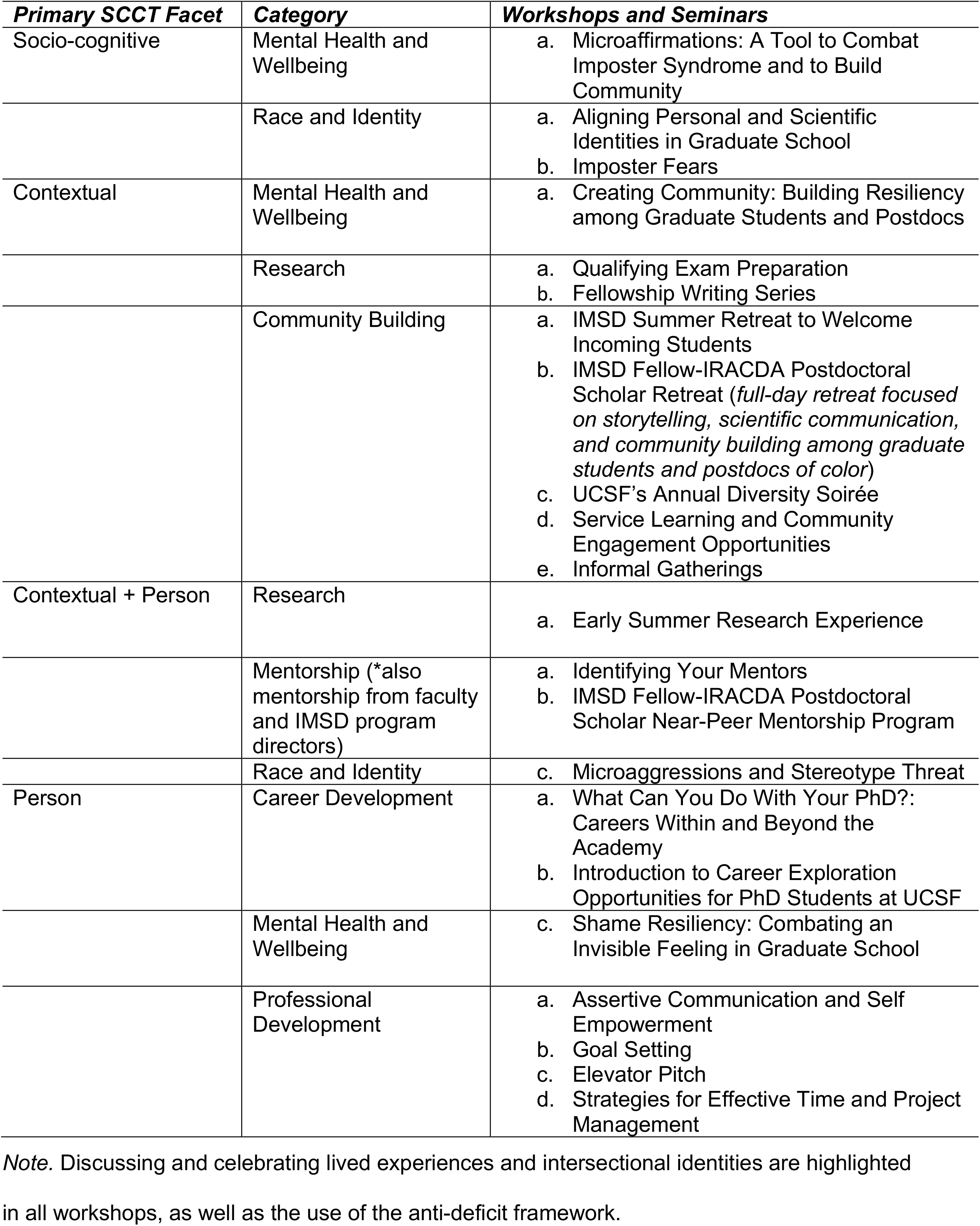
UCSF’s IMSD program components by social career cognitive theory facet

## Description of Findings and Their Implications

We interviewed 17 of 18 (94%) students at mid-program and 14 of 18 (78%) at end-of program for a total of 31 of 36 (86%) interviews. Themes from the mid- and end-program interviews were similar and hence we describe the final themes combined and articulated differences by program trajectory where relevant. To provide the overall context for our sample, we first describe overall fellow experiences and perceptions of the future. We then describe our themes and the relationships amongst them, further supporting figure 1’s description of how one can apply an anti-deficit framework for transformative practices.

### Overall Experience

Graduate students reported positive experiences with the IMSD activities at the mid- and endpoints of the program. Most students noted engagement with IMSD alumni (senior students) who championed for IMSD and the positive benefits the program had had on their success in graduate school. The students stated that the program provided a wide range of useful information and resources, as well as many opportunities they would not have otherwise known about to successfully progress through their graduate studies. Given the stress, workload, and transitions in the first year of graduate school, some fellows acknowledged the unique benefits of IMSD offering financial support, along with generous professional and social support, including facilitating easier and more accessible networking opportunities between IMSD students and faculty. For example, one student expressed deep gratitude for obtaining information about the graduate school experience and described the importance of resources specifically provided for racial and ethnic underrepresented students who often may not be as advantaged coming into graduate school with limited access to resources and mentorship than their non-underrepresented counterparts. Using SCCT as an evaluation tool, we are able to measure success through relationship building between IMSD fellows and alumni, faculty, and program leadership, and multiple programs aspects.

### Summer Rotation

Graduate students emphasized the benefits of having an additional opportunity to conduct summer research prior to starting graduate school and building early connections with other UCSF community members through IMSD, particularly in building confidence and assisting in their transition into graduate school. IMSD students appreciated that it allowed them to get acquainted with the campus without the additional pressure of simultaneously starting their courses. Some students selected their summer research rotation lab as the one they would ultimately join for their dissertation research, which is a complex decision that requires interpersonal and scientific alignment

### IMSD Program Workshops and Seminars

The workshops and seminars are designed using an anti-deficit framework, with a complementary focus on aligning students’ personal identities (who they are as people) and professional identities (who they are as scientists). The workshops and seminars offer students an opportunity to engage in: a) career and professional development enrichment and mentorship activities that complement academic research training; b) peer-to-peer learning; and c) an environment to showcase, celebrate, and value individual personalities, identities, and experiences; all of which contribute to enhancing IMSD students’ sense of self and cultivating relationships that ensure long-term academic success. Furthermore, these events also provided invaluable community building and networking opportunities with speakers, faculty, and postdoctoral scholars who are specifically invested in providing ongoing support for HUMs.

As they progressed through their first two years of graduate school, IMSD helped reinforce skills and information they had learned in their degree programs and lab experiences. Most of the IMSD students also valued sessions focused on navigating the graduate school experience as a HUM student, such as the stereotype threat workshop, the discussion of imposter syndrome in graduate school, and having the chance to openly share their personal, and sometimes vulnerable, experiences on the role of race and identity in science and mentorship, which is often not discussed regularly in their graduate programs. Another student expressed that they had gained knowledge of the types of opportunities and support available, where to seek assistance both within and outside of the institution. Through the evaluation process, we learned that we could better support students’ success through the inclusion of more individualized feedback, more career exploration opportunities, and specific types of research guidance.

### IMSD Community Building Activities

IMSD students valued the support from being in a community of students with shared experiences and benefited from branching out beyond their scientific or degree-based communities to cultivate additional sources of support. Some students suggested using group meetings as times for mentorship between 1^st^ year and 2^nd^ year students, as well as potentially formalizing mentoring relationships between current IMSD students and program alumni. This suggestion exemplifies some of the transformative practices that one can implement through program development and evaluation that our use of SCCT and AD facilitated.

In addition to the group monthly meetings, IMSD students appreciated less formal community building events and opportunities to interact with IMSD alumni and current students, which allowed for peer-to-peer mentorship through the sharing and learning from other students’ experiences and resources. As one example, students enjoyed having the opportunity to meet and support the incoming IMSD first-year cohort at the summer retreat. Social events that occurred earlier in the program were particularly helpful in building much needed social support, and several students asked for additional activities that focused on personal and professional guidance from program alumni, prior to starting graduate school. All students stated that they interacted with fellows in their cohort during and outside of the IMSD programming, and generally appreciated not feeling alone as HUMs in science.

### IMSD Mentorship Experiences

Most IMSD students also had positive bi-directional mentoring interactions, including those who leveraged the opportunity to establish relationships with IMSD alumni. IMSD alumni were described as approachable and genuinely invested in students’ academic success. IMSD alumni were especially helpful with social support, including, sharing how to navigate living in the San Francisco Bay Area, guiding students on future courses within their program, and providing ongoing information regarding possible career choices; all of which help current IMSD students make the most out of their experience in graduate school

In addition to the benefits of support from IMSD alumni, IMSD students described IMSD program leaders as a key source of guidance, valued their investment in students’ well-being and success, and appreciated their commitment to ongoing improvements in the program. Students recognized the difficulty faced by the program leaders in bringing the cohort together from different degree programs with varied schedules. Students described that consistent opportunities for interaction throughout the program with IMSD program leaders, who also served as mentors, were vital to their success as it signaled clear leadership and cohesion.

### Future Involvement with UCSF’s IMSD

Almost all students expressed an intent and enthusiasm to remain involved with UCSF’s IMSD in the future. They hoped to help future students and give back to the program which they felt “carried” them through graduate school. For example, a few students expressed an interest in teaching at the undergraduate level to inspire rising students and give back to the community, while others expressed a desire to be IMSD student faculty advisors in the future. All students shared an interest in mentoring incoming cohorts and participating in future IMSD events such as workshops or panels. As one student described, they would want to continue to be mentored by IMSD students and program leadership. These findings emphasize the importance of relationship building for success.

## CONCLUSION

Diversity, equity, and inclusion practitioners need to apply an anti-deficit framework to quickly adapt to the evolving goals, interests, and capabilities of STEM PhD students in a manner that is proactive about their retention, achievement, and success. Previous research on the experiences of HUM students in biomedical sciences using SCCT has suggested that institutional leadership needs to go beyond individual skill-building to attend to individual values and the promotion of systemic reforms to better achieve the intended outcomes of increasing diversity in student acceptance and enrollment in these types of graduate programs (Gibbs & Griffin, 2013; Maher et al., 2017; St. Clair et al., 2017). Such calls for equity-minded reforms are progressive and needed. However, without clearly stipulating the lens through which individual values and systemic reforms should be identified and promoted, practitioners run the risk of missing nuanced dynamics at play and unintentionally perpetuating persistent inequities in their institution (Malcom-Piqueux & Bensimon, 2017). Professional development strategies, even those that attend to personal values and institutional relationships, may define and measure success by the individual’s willingness or ability to conform to pre-existing structures or superimposed ideas (Malcom-Piqueux & Bensimon, 2017). Innovative programming requires practitioners to have a shared perspective and methodology to challenging these systemically engrained practices. The anti-deficit framework facilitates this by refining calls to action through the centering and uplifting of students’ identities as a mechanism to elucidate the complex relationship between individual’s and institution’s norms, values, and practices. This perspective orients program development and evaluation around supporting student retention and flourishing in a sustainable educational environment that values personal and professional relationship building.

We chose to complement our use of anti-deficit framework with SCCT because those facets work best with our IMSD programming; however, this approach can work with other theories as it relates to program goals. Other institutions whose work may not be explicitly guided by SCCT can also benefit from evaluating and refining their programs through an anti-deficit framework by referring to figure 1 and exchanging SCCT for their theory of choice. To adopt this approach, one would revisit the underlying research questions and assumptions of their work to ask, *“Is this approach accounting for the lack or mismatch within the student, or is it attending to the enablers of success?”*. By iteratively identifying and emphasizing the enablers of success and facilitating structured community and social support for personal and professional identity development, institutions can create, redesign, and sustain programming that supports increased academic and scientific development for HUM PhD learners.

## Footnotes

1 Some diversity literature uses the term “underrepresented minority (URM)” as it matches the institutional language being used in the literature and funding mechanisms supporting diversity, equity, and inclusion work. However, we the researchers and the students we interact with prefer more contemporary terminology such as “historically/presently marginalized populations” or “historically underrepresented and marginalized students” which more explicitly address the complex structures and power dynamics at play.

2 Historically underrepresented and marginalized students include U.S. citizens or permanent residents who are African American, Hispanic American, Native American, and natives of the U.S. Pacific Islands, and individuals living with disabilities.

## Notes

We have no known conflict of interest to disclose. The National Institute for General Medical Sciences (NIGMS) Initiative for Maximizing Student Development (IMSD) fellowship is funded by grant award 5R25GM056847.

### Competing Interest Statement

The authors have declared no competing interest.

